# Functional memory of drought affects leaf chemical defenses and microbial interactions in aspen

**DOI:** 10.1101/2025.11.17.688914

**Authors:** Aubrey M. Hawks, Jaycie C. Fickle, Kelly L. Kerr, Dale L. Forrister, Allison M. Perkins, Efthymia Symeonidi, Madelyn Allen, Cannon Fehrenbach, Hsiao-Nung Chen, Richard L. Lindroth, William R.L. Anderegg, Talia L. Karasov

**Author notes:** Contributed equally (William R.L. Anderegg, Talia L. Karasov).

## Abstract

Drought is an increasingly important driver of tree mortality, but its long-term impacts on trait expression remain poorly understood. Leaves are a key interface at which trees respond to abiotic stress, and foliar traits such as chemical defenses and microbial communities may play an important role in determining post-drought resilience. Here, we tested whether prior-year drought leaves a persistent imprint on leaf traits in *Populus tremuloides.* Using a three-year common garden experiment with controlled water limitation, we measured two major classes of defensive phenolics—salicinoid phenolic glycosides (SPGs) and condensed tannins (CTs)—alongside foliar fungal community composition. SPG concentrations increased and CTs declined in response to prior-year drought, with these effects persisting across growing seasons. While overall fungal community composition remained relatively stable, we detected shifts in the relative abundance of individual amplicon sequence variants (ASVs), particularly within potentially pathogenic lineages, associated with drought history. These findings provide evidence that drought leaves a legacy in leaf chemistry and microbial colonization, with potential consequences for how trees tolerate future abiotic and biotic stress. Our results highlight the importance of incorporating foliar trait legacies into models of forest resilience under climate change.

**Significance Statement:** Trees are known to exhibit long-term responses to drought, but whether such legacy effects extend to annually renewed tissues like leaves remains unclear. Our findings reveal that prior-year drought alters both chemical defenses and microbial associations in newly formed leaves, demonstrating a form of physiological memory. This work uncovers a novel mechanism by which trees integrate past environmental stress into future interactions, with implications for understanding forest resilience under climate change.

## Introduction

Hotter, drier, and more prolonged droughts linked to climate change are now recognized as a dominant driver of tree mortality, with impacts that can extend for years after drought conditions subside (1–3). Despite this recognition, current models remain poor at predicting when and where drought will trigger widespread die-off (4, 5). Two factors are increasingly recognized as central to this limitation: high individual-level variation, such as differences in past stress exposure or microclimate, and interactions with biotic agents, including pests and pathogens (4, 6, 7). Biotic agents can intensify—or in some cases reverse—the expected effects of drought depending on the identity of the agent, the host species, and the severity of stress (8–11). These dynamics are insufficiently represented in current models, largely because we lack mechanistic understanding of how drought alters tree–biotic interactions and how those interactions amplify or dampen mortality risk (12). Because foliar traits mediate interactions with both abiotic stressors and biotic agents, they may be critical for understanding drought-driven mortality. Among these, defensive chemistry and foliar microbial communities stand out as traits that directly influence how trees interact with pests, pathogens, and stress (13, 14). Key unknowns are how drought reshapes leaf chemistry and microbial communities, how persistent these changes are, and whether interactions between the two contribute to tree resilience or vulnerability.

Leaves are a critical interface where trees encounter both abiotic stress and biotic attack, and traits expressed there are likely to mediate how drought influences tree performance. Plants invest substantial resources in specialized metabolites—up to 5% of their photosynthate and as much as 20% of available nitrogen (15–17)—despite the associated costs to growth, because these compounds improve stress tolerance and defense (18, 19). By influencing herbivores, microbial colonization, litter decomposition, and nutrient cycling, these compounds generate cascading ecological effects. Many of the most abundant plant metabolites are antioxidants that mitigate abiotic stress directly by scavenging reactive oxygen species during drought (13, 20, 21). Foliar microbes, in turn, can shape tree health by enhancing stress tolerance or, conversely, by acting as opportunistic pathogens (14, 22, 23). Their composition reflects both environmental filtering and host traits, including chemical defenses, positioning chemistry and microbes as potentially intersecting mediators of drought outcomes and community composition(24).

Evidence that drought influences allocation to specialized metabolites comes from a wide range of studies, with stronger experimental support in crop species than in forest trees. Shifts in terpenes and phenolics have been measured following short periods of drought, though the magnitude and direction of responses vary with drought severity and plant species (20, 25–34). In conifers, such shifts can have direct ecological consequences; for example, decreases in monoterpene concentrations in *Pinus edulis* have been associated with increased bark beetle attack during drought (11, 35, 36). More recent research suggests the potential for longer term impacts of severe drought stress. Laoué et al. (2024) found reduced flavanol allocation in *Quercus pubescens* that persisted for up to two years, and Eisenring et al. (2024) documented lasting metabolomic shifts in *Fagus sylvatica* following drought-induced defoliation (37, 38). These studies, conducted under field and semi-natural conditions, indicate that drought may leave enduring legacies in defensive chemistry. Replicated and randomized experiments are needed to assess whether drought consistently reshapes defensive chemistry, to clarify the mechanisms driving these responses, and to determine if such changes translate into altered biotic interactions.

Evidence that drought alters plant-associated microbial communities is accumulating rapidly, and water availability is emerging as a major determinant of the leaf microbiome (39–41). These effects are less well documented in forest trees, though rainfall-exclusion experiments in *Quercus ilex* have reported increased bacterial and fungal diversity in the phyllosphere following drought (42, 43). In *Populus deltoides*, drought altered microbial communities in a sex-dependent manner, with female trees hosting assemblages more resilient to pathogen infection (44). Shifts have also been reported in conifers, where changes in bacterial communities were observed under rain exclusion, though this response may reflect altered inoculation rather than direct stress-mediated effects (45). Longer-term drought effects on forest microbial communities have been documented in soils, but their persistence in foliage is much less certain (46, 47). Moreover, the links between drought, foliar chemistry, and microbial colonization remain poorly understood.

We examined how drought history shapes specialized metabolism and foliar microbial communities post-drought using quaking aspen (*Populus tremuloides*). This widely distributed and chemically well-characterized tree species serves as a model for studying plant defense and plant–microbe interactions in forest ecosystems. Aspen produce two major classes of specialized metabolites—salicinoid phenolic glycosides (SPGs) and condensed tannins (CTs)—which are involved in both biotic and abiotic interactions. SPGs are salicylate derivatives largely restricted to the Salicaceae (e.g., *Populus* and *Salix*), while CTs (also known as proanthocyanidins) are flavan-3-ol polymers found broadly across plant taxa (48)). In aspen, allocation to SPGs and CTs varies widely, and each can range from less than 1% to over 30% of leaf dry weight. Both are products of the phenylpropanoid pathway, and their concentrations are often inversely correlated, suggesting a resource-based trade-off driven by shared biosynthetic precursors. SPGs are strongly associated with herbivore deterrence: for example, each 1% increase in SPG leaf dry weight reduces female spongy moth larval weight by approximately 73.9 mg (0.5%) and extends development time by 1.1 days (49). Though they have been shown to reduce food preference in chrysomelid beetles, CTs are thought to have limited effects on herbivores and are more often associated with anti-microbial activity (48, 50). Experimental work has shown that CTs can inhibit *Melamspora larici-populina*, a biotrophic rust fungus infecting *Populus*; at 0.5 mg/mL they reduced spore germination by 4.2-fold in vitro (51). Natural surveys of foliar communities in *Populus* further suggest that microbial assemblages are shaped by environmental conditions, with drier sites often harboring distinct community profiles (52).

We utilized a three-year common garden drought experiment established by Kerr et al. to test whether these traits capture drought legacies in a controlled setting (Figure 1) (53, 54). From 2021 to 2023, two blocks each were maintained under consistent control (CCC) or drought (DDD) conditions. An additional three blocks alternated treatments to simulate shorter drought periods (CDD) or recovery years between droughts (DCD) (Figure 1C). We sampled leaves at peak drought each year to address three questions: (**1**) **Does drought alter the chemical profile of aspen leaves, and how long do these effects persist?** (**2**) **Are foliar microbial community constituents distinct in trees that experienced prior drought? and** (**3**) **Do concentrations of SPGs or CTs correlate with shifts in foliar microbial community composition?**

**Figure 1.**
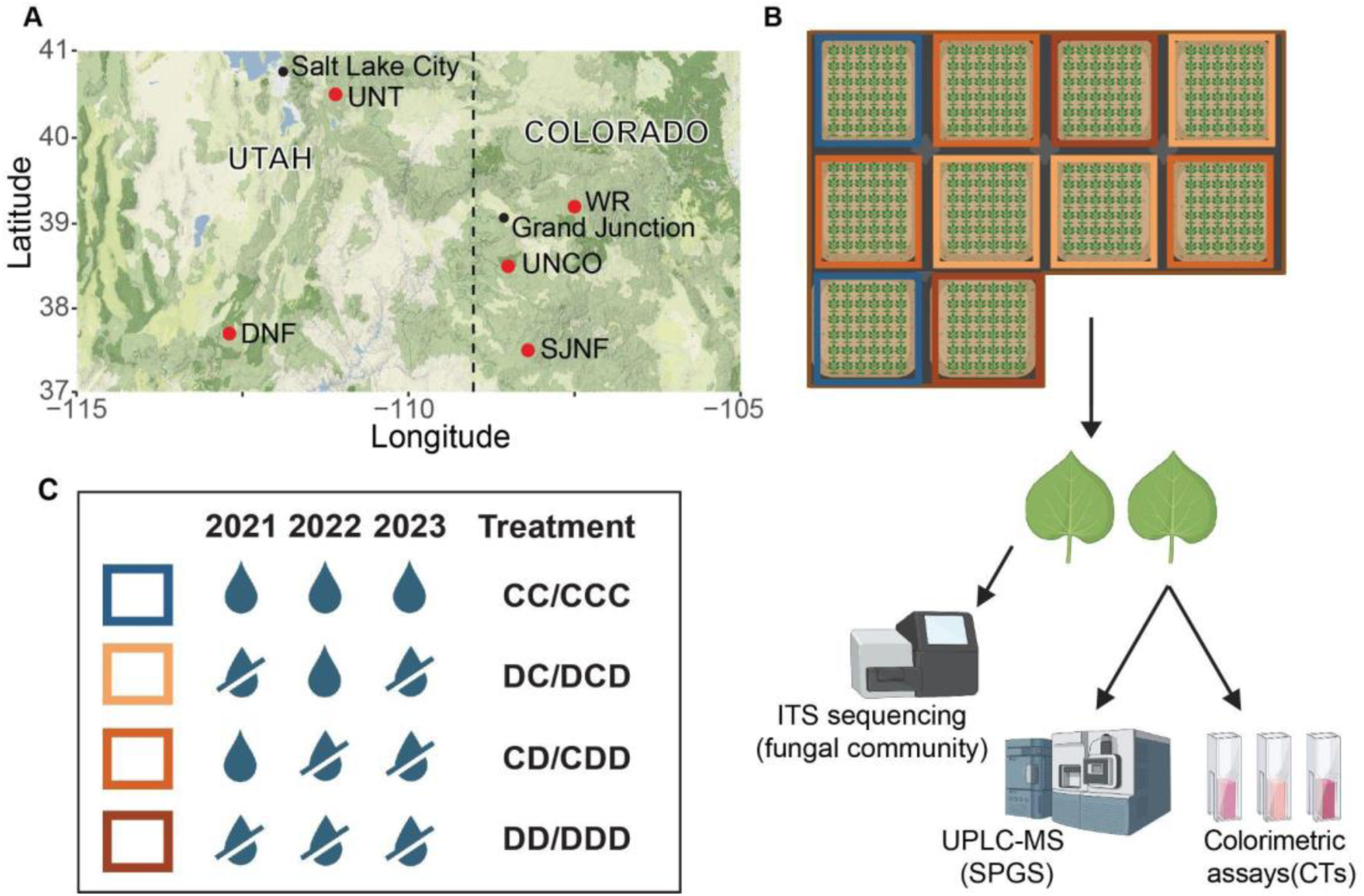
Experimental setup for common garden. Field sites were established at locations across Utah and Colorado (A). Root stock taken from multiple stands at each site was propagated and planted into randomized common garden blocks (B). From 2021–2023, trees in the common garden received either normal watering or drought stress from June to September. Water icons indicate treatments with colors and abbreviations corresponding to treatment groups in subsequent figures (2022 treatment/2023 treatment) (C). Samples were collected for analysis of specialized metabolites and microbial communities (B).

## Results

### Prior year drought affects production of SPGs and CTs

We measured the full suite of foliar metabolites using Ultra Performance Liquid Chromatography – Mass Spectrometry (UPLC-MS), and determined absolute concentrations for the focal SPG class using purified salicin, salicortin and tremulacin standards. We first used dimensionality reduction to determine the major sources of variation in chemical profiles between samples using all identified compounds. In 2022, PCoA1 of the UPLC data captured 67.8% of variation, with the majority of contributors corresponding to peaks for the four major SPGs: salicin, salicortin, tremulacin and tremuloidin (supplementary figure 1A). PCoA1 from 2023 (68.1% of variance) was also dominated by SPGs(supplementary figure 1B). While there was no significant separation between treatment groups in 2022, in 2023, trees experiencing two or three consecutive years of drought (CDD, DDD) separated from those with alternating drought and normal watering (DCD, CCC) (ps = 0.002 vs CCC, 0.24 and 0.002 vs DCD for CDD and DDD respectively). DCD also separated from CCC (*p* = 0.24). PC2 described an additional 8.4% (2022) and 6.7% (2023) of variance and was associated with larger, more polar compounds that eluted early in the gradient. These patterns are consistent with prior literature that indicates that SPGs are a major component of chemical variation in aspen and suggests that they may vary according to treatment groups.

Linear mixed-effect models controlling for effects of genotype indicated that there was no significant difference in total SPGs between trees experiencing current-year drought and those with normal watering in 2022 *(p* = 0.3417) (figure 2A). Trees that experienced drought the prior year had higher SPG content by 1.46% dry weight, an 11% increase over plants that were normally watered the prior year (p <0.0001), driven by increases in salicortin and tremulacin in the drought-control and drought-drought treated trees (supplementary figure 2). Prior year water availability explains 7.7% of variance In total SPGs between samples, compared to 14.5% explained by genotype(Table 1). CT concentrations, as measured by colorimetric acid-butanol assays, did not differ between treatments in 2022(figure 2C).

**Figure 2.**
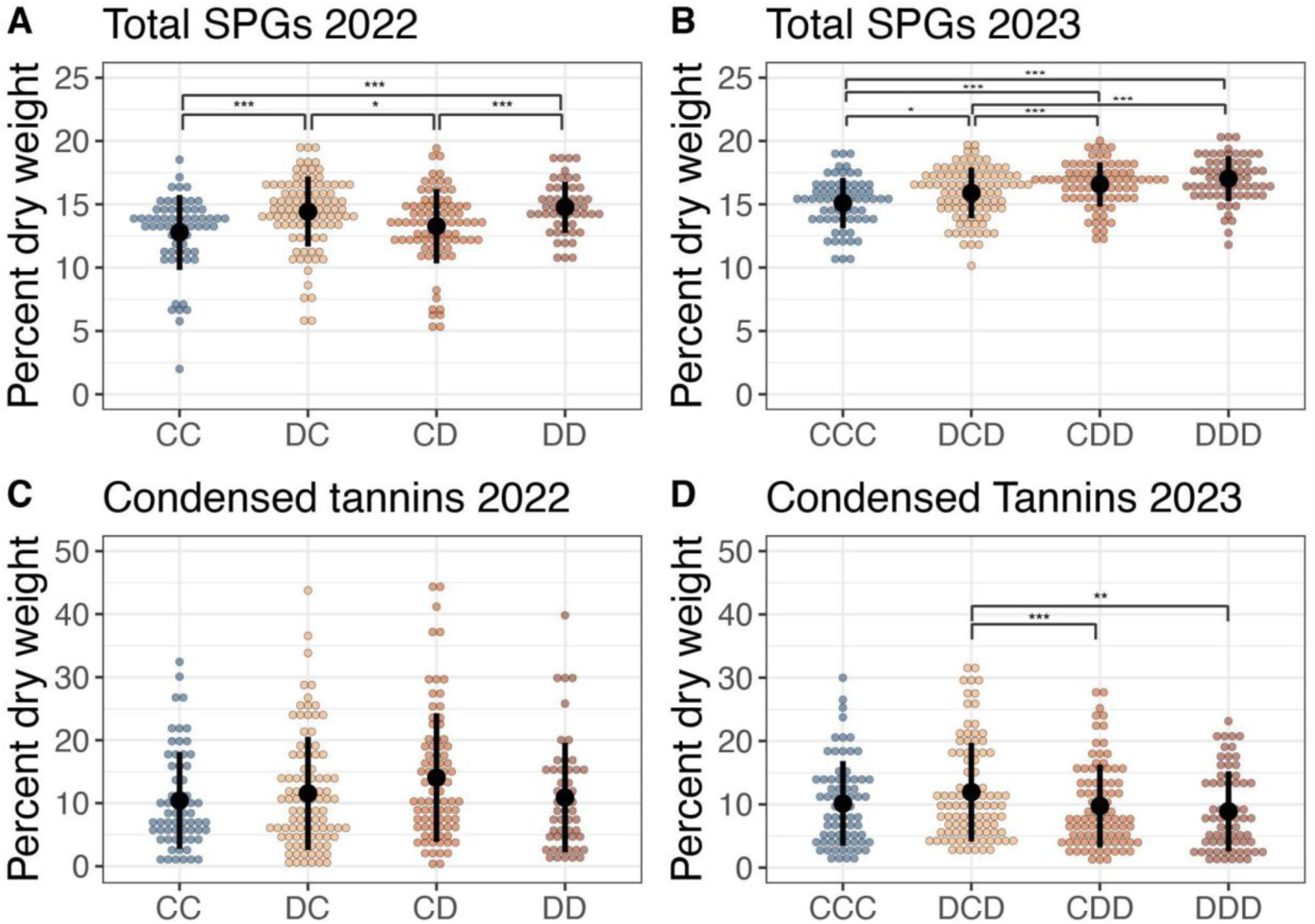
Memory of drought shapes aspen allocation to foliar defensive chemistry. Allocation to SPGs was higher in trees that experienced drought in the prior year in both 2022 (A) and 2023 (B). CTs did not differ between treatment groups in 2022 (C), but were significantly lower in previously drought-stressed trees in 2023 (D). Colored dots represent individual samples; black lines show means ± SE. Pairwise comparisons were performed using *emmeans* with Tukey-adjusted p-values: (***) < 0.001, (**) <0.01, (*) <0.05. Sample sizes: n ≤ 51 (CCC, DDD) or n ≤ 74 (CDD, DCD).

**Table 1.** Variance explained (VE) in chemical defenses for treatment and genotype.

|  | year | VE by genotype (%) | VE by treatment (%) | CI lower | CI upper |
| --- | --- | --- | --- | --- | --- |
| Salicin | 2022 | 1.2 | 9.3 | 6.6 | 14.1 |
|  | 2023 | 20.5 | 35.5 | 29.9 | 41.0 |
| Salicortin | 2022 | 15.3 | 8.9 | 3.3 | 13.0 |
|  | 2023 | 38.1 | 10.7 | 6.9 | 17.1 |
| Tremulacin | 2022 | 17.3 | 6.3 | 3.2 | 13.6 |
|  | 2023 | 56.5 | 5.5 | 4.6 | 8.2 |
| Total SPGs | 2022 | 14.5 | 7.7 | 5.1 | 11.8 |
|  | 2023 | 46.3 | 10.9 | 8.4 | 15.0 |
| CTs | 2022 | 40.8 | 1.3 | 0.9 | 3.9 |
|  | 2023 | 50.8 | 3.1 | 1.3 | 3.6 |

### Prior year effects still dominate after prolonged drought

In 2023, total SPGs again differed primarily by prior year treatment, with an 8% increase in plants that had experienced drought the previous year compared to prior year controls (*p* <0.0001) (figure 2B). DCD plots were also significantly higher than controls (5%, *p* = 0.0173), despite sharing the same prior year treatment, a difference driven by significant increases in both salicin (26%, *p* <.0001) and tremulacin (4%, *p* = 0.0254) (supplementary figure 2). It cannot be determined from the data whether this is due to the drought in 2021 or 2023 as these are correlated in the treatments. No significant differences were observed between the two and three year drought groups. Both genotype and treatment explained more variance in 2023 (46.3 and 10.9%, respectively), with treatment effects strongest for salicin and salicortin (Table 1). CTs in 2023 were significantly higher in the alternating drought treatment than in the two (31%, *p* = 0.0006) and three year (32%, *p* = 0.0011) consecutive drought plots (figure 2D).

### Evidence for a shift in allocation between SPGs and CTs following drought

Comparisons among treatments revealed contrasting allocation patterns between SPGs and CTs in plots that switched treatments between 2021 and 2022. Between 2022 and 2023, all groups showed modest SPG increases, but the increase in CDD was 2.93 fold greater than in DCD *(p*<0.0001) (figure 3A, supplementary figure 3). In contrast, CT concentrations increased in DCD but decreased in CDD by a mean of 6.095% dry weight, with the year-to-year change differing by a mean of 6.77% dry weight between the groups (p = 0.0014) (figure 3B, supplementary figures 3-4).

**Figure 3.**
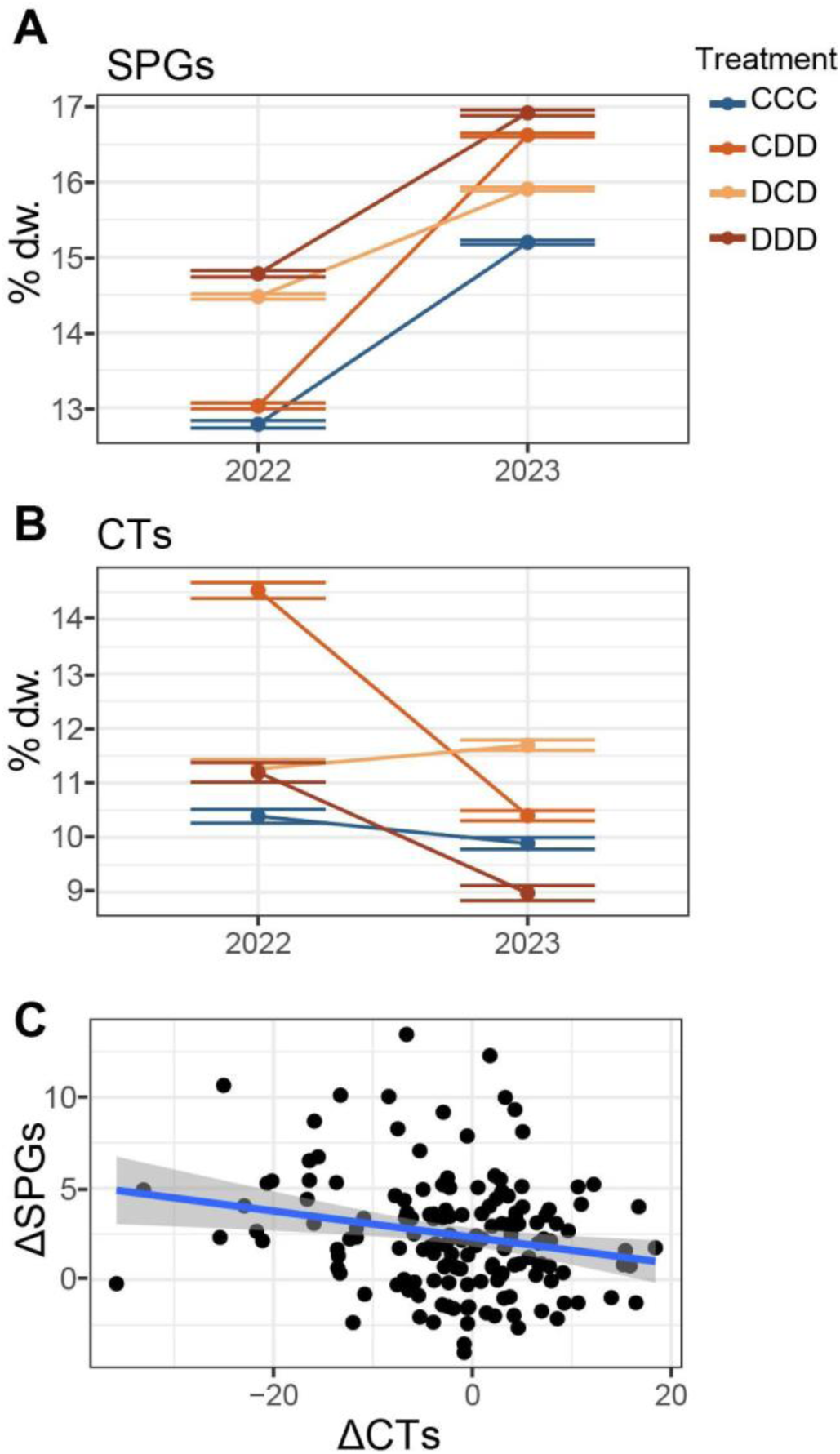
Drought legacy-induced SPG increase is associated with decrease in CTs. Mean total percent dry weight ± SE for SPGs (A) and CTs (B) in 2022 and 2023 by treatment group. In the CDD and DCD groups, changes in SPGs from 2022 to 2023 were inversely correlated with changes in CTs (slope = –0.0718, adjusted R² = 0.040, *p* = 0.0074).

Consistent with prior literature, total SPG content and CT content were negatively correlated (slope = –1.16, adjusted R^2^ = 0.1084, *p* <0.0001). To determine whether individual trees were shifting their allocations between these two compound classes, we tested for correlation between year-to-year changes in SPGs and CTs within the groups that switched treatments between 2021 and 2022 (CDD and DCD). Changes were negatively correlated (slope = –0.07181, adjusted R^2^=0.03996, *p* = 0.0074) (Figure 3C), supporting the hypothesis of an allocation trade-off between SPGs and CTs due to drought induction.

### Potential pathogens show association with prior year treatment

Our survey of foliar fungal communities at peak drought identified 6,467 unique ASVs in 2022 and 8,290 ASVs in 2023 after filtering for family-level assignments, with *Cladosporium* and *Comoclathris* the most abundant taxa across treatments—consistent with previous *Populus* phyllosphere surveys (figure 4a, supplementary figure 5) ((52). We compared community composition in the common garden in 2022 with that of the field sites and found that the garden composition broadly corresponded to composition in the field, with no significant differences in colonization by *Cladosporium*, *Didymella*, *Dothidotthia*, or *Fusarium* (Benjamini Hochberg (BH)-adjusted *p* > 0.01). Field sites differed primarily in the abundance of several potentially pathogenic genera (*Endoconidioma*, *Cytosporia*, *Gogovinomyces*, *Septorellia*; BH-adjusted *p* < 0.0001), driven mainly by expansions at individual sites. Garden samples had higher *Comoclathris* abundance (BH-adjusted *p* < 0.0001) and lower overall diversity than field samples (Shannon diversity difference = –0.456, *p* < 0.0001), indicating that the common garden is a reasonable but slightly less diverse representation of microbial communities associated with these genotypes (figure 4b).

**Figure 4.**
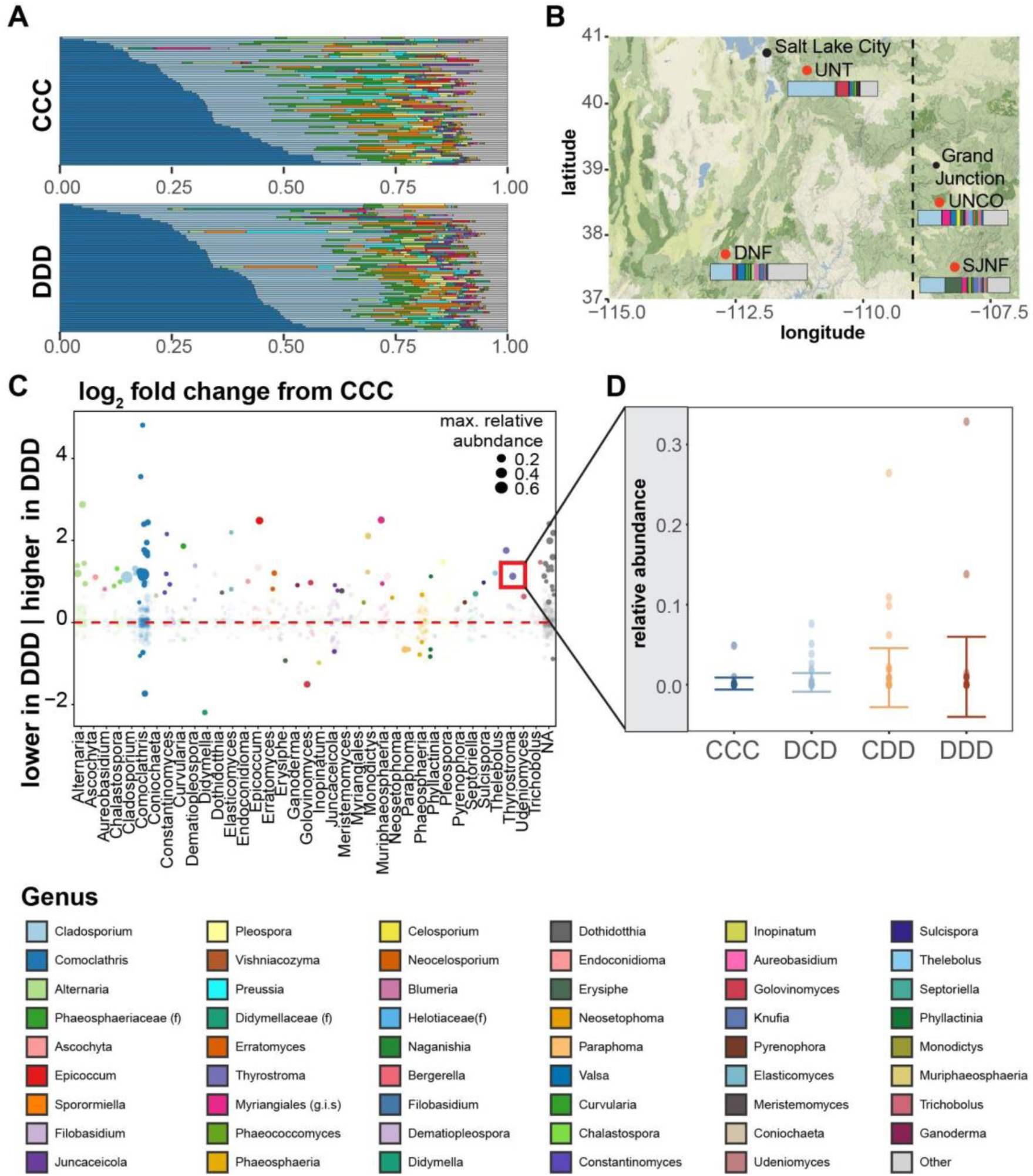
Abundances of potentially pathogenic fungal ASVs differ according to two-year drought history. Samples collected at peak drought in 2023 (garden) and 2022 (field). ITS1 amplified from single leaves collected in the common garden. Relative abundances of fungal genera in fully watered trees (CCC) and those experiencing prolonged drought (DDD) (A). Mean community composition from field-site leaves showed more diverse fungal communities but similar constituents to common garden trees (B). Numerous potentially pathogenic lineages differed in prior-year drought-stressed trees (C); points represent individual ASVs scaled by maximum relative abundance across samples. Points below the dashed red line were more abundant in normally watered trees; points above were more abundant under the contrasted drought regime. Opaque points indicate an adjusted p < 0.05. Relative abundance in individual samples across all treatments for a single ASV (*Thyostroma*) found to be at higher abundance in DDD vs. CCC (D).

Drought was associated with increases in the *Dothidiomycetes* class in both 2022 and 2023 (BH-adjusted *p* < 0.01 in 2022; *p* <0.0001 in 2023) (Table 2). In 2023, this included significant increases in *Myriangiales* and *Didymellaceae* (BH-adjusted *p* < 0.0001 and <0.05, respectively). Drought was also associated with an increase in *Corticiales* (BH-adjusted *p* <0.05) and a decrease in *Erysiphales* (BH-adjusted *p* < 0.0001) in 2023. Prior-year drought showed no detectable effects on community composition down to the genus level in either year.

**Table 2.**
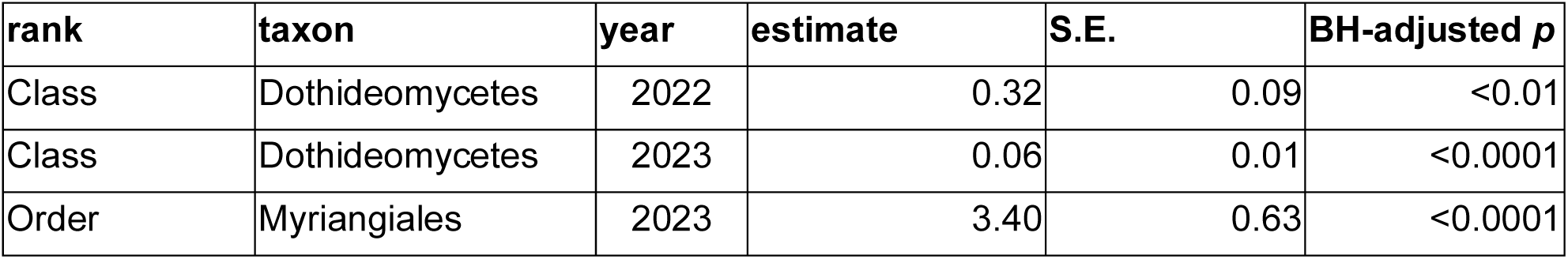
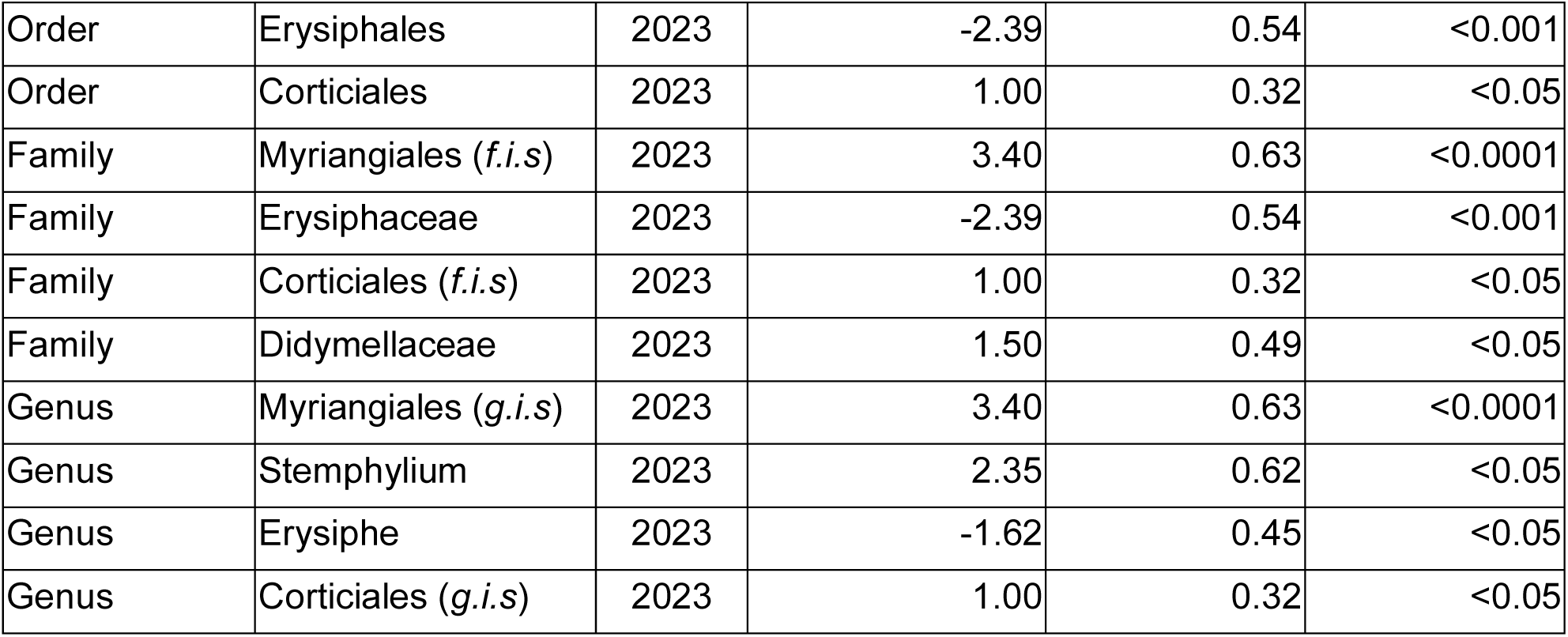
Linear mixed-effects model results for drought effects on fungal taxa.

To better understand how drought history shapes fungal diversity at finer taxonomic scales, we next looked at ASV-level variation in abundance. In order to better assess whether drought has an impact on pathogen infection, we restricted our analysis to families with known or suspected plant pathogens and further filtered for ASVs with variance in relative abundance in the top quartile between samples. We identified 4,152 potentially pathogenic ASVs in 2022 and 4,272 in 2023. We then performed differential abundance testing, contrasting each drought treatment (DC/DCD, CD/CDD and DD/DDD) with the control group (CC/CCC) using the likelihood ratio test in deseq2, with garden plot as a covariate in the model to control for spatial structure(55).To examine whether prolonged drought affected colonization by potential pathogens, we focused on the contrast CCC vs. DDD contrast. We found 81 ASVs that were more abundant under prolonged drought (DDD), 45 of which were not significantly enriched in either of the intermediate drought treatments (DCD,CDD; *p* <0.05). Of the 15 ASVs that were less abundant in DDD compared to CCC, 4 were unique to three years of drought.

To explicitly test whether prior year drought influenced the ASV composition--while avoiding confounding effects of current year drought--we compared CC and DC from samples from 2022. The analysis identified 49 ASVs enriched under prior-year drought and 15 enriched in prior-year controls, after accounting for effects of garden plot. These results suggest that legacy drought can influence colonization success of specific fungal lineages even when broader community composition remains unchanged.

Finally, we used linear mixed effect modeling to establish whether defensive chemistry was associated with fungal community shifts. *Sporormiaceae* abundance was positively associated with SPG concentrations (BH-adjusted *p* < 0.05). No significant associations were detected between quantities of CTs and any fungal taxa down to the genus level.

## Discussion

Specialized metabolites are key determinants of tree tolerance to biotic and abiotic stresses. Genetic variation is the principal determinant of SPG and CT production in aspen grown in our common garden in Utah. However, our findings demonstrate that the foliar chemistry of aspen does exhibit memory of past environmental stress: prior year drought had the most pronounced effect on both compound classes. Responses varied widely—some genotypes exhibited sharp switches between SPG– and CT-dominated profiles, while others changed little—suggesting that drought legacy effects may unevenly influence plant–biotic interactions across populations.

SPG allocation is generally stable throughout the growing season and only minimally inducible (56, 57). In our study, SPG concentrations increased following prior-year drought, consistent with a response to environmental cues experienced before bud break or during the preceding year rather than to immediate water stress. SPGs showed no detectable response to current-year drought, but prior-year effects were evident in both 2022 and 2023, and of similar magnitude across years. The persistent effect on SPGs suggests that a single drought event can leave a lasting chemical signature into the following year, regardless of subsequent conditions.

CTs are typically more inducible than SPGs, responding to herbivory, nutrient limitation, and light (57–59). Contrary to expectations, we found no short-term CT increase following drought; instead, CTs decreased as SPGs increased, consistent with a trade-off in phenolic allocation. CT concentrations were unaffected by current-year drought, and no significant differences were detected in the second year under either current or prior drought conditions. The only clear effect emerged in the third year (2023). This pattern may reflect the weaker, less uniform drought measured in 2021, which likely produced only subtle changes in CTs in 2022, although similar trends were apparent. Alternatively, multiple consecutive drought years may be required to elicit a strong CT response. The high variability among genotypes in our dataset suggests that some individuals may be more chemically flexible than others, with potential implications for differential resilience to future stresses.

The trade-off between SPGs and CTs has been consistently demonstrated in both surveys of variation within genotypes and through experimental manipulation of expression in the phenylpropanoid pathway (19, 48, 56, 60, 61). Evidence for a trade-off due to environmental induction is mixed. Here, our data suggest that such constraints may partially shape drought responses—but with substantial inter-genotypic variation in magnitude and direction.

Even modest shifts in the SPG–CT balance can carry functional significance. CTs contribute to abiotic stress tolerance—mitigating photosystem damage and oxidative stress under drought and UV-B in *Populus* hybrids (62, 63)—and defend against fungal pathogens like *Melampsora* spp., while influencing microbial diversity and decomposition(64, 65). Higher SPG concentrations have consistently been associated with decreases in herbivore fitness for several major pests of aspen (49, 66–68). They may also confer UV protection. Individual-level, drought-linked chemical shifts—as seen in our data—could thus meaningfully alter herbivore performance, pathogen pressure, and stress resilience.

Taken together, these results show that prior-year drought increased SPGs, decreased CTs, and altered the relative abundance of a subset of potentially pathogenic fungal strains though a causal connection between the two has not been established. Many major foliar pathogens of aspen, including *Melampsora* rusts and *Venturia* leaf blight, are more prevalent under well-watered conditions, while others, such as *Cytospora chrysosperma* and *Septoria* spp., are opportunists that exploit stressed or wounded hosts (69–71). Some are insect-vectored or invade through herbivore damage. Although we lacked herbivore data, the magnitude of SPG and CT changes we observed has been shown in previous studies to affect herbivore performance and, by extension, likely pathogen incidence. These drought legacies may produce heterogeneous outcomes for both herbivore and pathogen pressure in subsequent years.

In deciduous trees like aspen, the physiological and molecular mechanisms behind drought stress memory remain poorly understood. In woody tissues, drought induced damage to fine roots and xylem can limit water and nutrient uptake long after the period of drought, reducing resource availability and potentially constraining how leaves allocate resources to metabolism. Alternatively, persistent changes in carbohydrate reserves due to reduced photosynthesis during drought or alterations to regulation of specialized metabolite pathways could produce altered chemical profiles following drought stress. In other plant systems, drought-induced epigenetic modifications, small RNAs and transcriptional reprogramming have been associated with changes to stress-tolerance following drought events, though few of these studies have been conducted in perennial plants or over multiple growing seasons(72–74). The paucity of documented examples in deciduous trees highlights a key gap in our understanding. Our results show a foliar chemical legacy of drought exposure, but further research is needed to ascertain whether the observed pattern is driven by active regulation or an indirect effect of structural damage to other plant tissues. It further remains to be determined how these changes interact with subsequent biotic and abiotic challenges and whether they are beneficial to post-drought recovery or likely to exacerbate the detrimental effects of drought.

In either case, these legacy effects may have far-reaching ecological consequences. In aspen, severe multi-year drought has been identified as the major driver of “Sudden Aspen Decline” (SAD) ((75, 76). Canopy dieback and mortality are often followed by increased prevalence of opportunistic pathogens such as *Cytospora chrysosperma* and insect pests(70). Our results suggest that drought-induced shifts in chemical defenses could be a contributing factor to post-drought vulnerability in aspen by altering host-pathogen or host-herbivore dynamics beyond the immediate period of drought. More broadly, climate change is predicted to intensify and prolong periods of drought in many regions(77, 78). Similar dynamics have been observed in other systems, for example, drought constrains induction of terpenes and increases susceptibility of *Pinus edulis* to bark beetle attack(11). This raises the possibility of more frequent large-scale stress-associated pathogen or pest outbreaks in forest trees. Understanding how drought legacies in plant chemistry mediate biotic interactions will be critical for predicting forest recovery and informing management strategies in a rapidly changing climate.

## Materials and Methods

### Field Site and Experimental Design

#### Field Site

Details of field site selection and establishment are provided in Kerr et al. (2023) ((53, 54). Briefly, sites were chosen to span a gradient from hot-dry to cool-wet climates, with similar elevation. Clonal genotypes from each site were verified using PCR amplification of eight microsatellite markers.

#### Common Garden

The common garden was established by Kerr et al. (53). Clones for the common garden were derived from field-collected rootstock in 2019. Rootstock was collected from individual stands at each field site and treated with 0.3% indole-3-butyric acid (Hormex Rooting Powder No. 3, Westlake, CA, USA) to promote root development. Once established, rooted cuttings were transferred to pots in a greenhouse and later transplanted outdoors in May 2020 into a common garden on the University of Utah campus. The garden comprised 10 randomized blocks, each containing a 6 × 6 grid of plants spaced 50 cm apart, totaling 360 individuals.

In 2021, garden beds were randomly assigned to control or drought treatments. All beds were watered to field capacity until mid-June. Drought treatments then received reduced irrigation—first to 50%, then to 25% of control levels—based on soil water measurements. Rainfall was not excluded in 2021. In 2022, two beds per treatment retained their original assignments, while three were switched from the previous year treatment (drought to control, and vice versa). In 2023, the two continuously watered control beds remained unchanged, while all others received drought treatment. Rain exclusion was implemented in 2022 and 2023 using overhead canopies installed before major precipitation events to intensify drought.

Drought severity was monitored via pre-dawn (4:00-6:00) and midday (12:00-14:00) leaf water potentials collected on a weekly basis throughout the drought treatments. A subset of 6 plants per bed were measured each week using a pressure chamber (Model 1505D Pressure Chamber Instrument, PMS Instrument Company, Albany Oregon, USA).

### Chemical Quantification

#### Leaf Sampling and Processing

Leaves were collected from the common garden in late August 2022 and 2023. One to two mature leaves per plant were clipped from mid-canopy height, placed in paper envelopes, and immediately stored on silica gel. All sampling was completed within 90 minutes at ambient temperatures below 30°C. Leaves were shaded during collection. Samples were dried on silica gel in the dark at 25°C for 48 hours, followed by vacuum centrifugation at 30°C for ≥12 hours. Dried samples were stored on silica gel in airtight bags at –20°C until analysis.

Leaves were powdered with glass beads in a FastPrep system at 4 m/s for 20 seconds. Approximately 15 mg of ground tissue was weighed for quantification of salicinoid phenolic glycosides (SPGs) and condensed tannins (CTs).

#### SPG Quantification

SPGs were extracted in methanol containing 0.01% β-resorcylic acid and quantified via UPLC-MS following Rubert-Nason et al. (2017) ((66). Analytical standards for salicortin and tremulacin were provided by R. Lindroth and purified as in Rubert-Nason et al. (2018) ((79). Salicin analytical standard was obtained commercially (≥ 99% purity by GC, CAS 138-52-3, MW 286.28 g/mol; Sigma-Aldrich, Cat. No. S0625). Peaks were detected and integrated using MZmine v4.3 via mzWizard with the following parameters: noise threshold (MS1) = 1E3, minimum feature height = 3.0E3, tolerance (scan-to-scan) = 0.02 m/z, tolerance (intra-sample) = 0.015 m/z, tolerance (sample-to-sample) = 0.02 m/z, smoothing and stable ionization across samples, retention time = 0.03-22.50 min, max peaks in chromatogram = 15, minimum consecutive scans = 4, approximate feature FWHM = 0.05 minutes, RT tolerance (intra-sample) = 0.2 minutes, RT tolerance (sample-to-sample) = 0.2 minutes(80).Duplicate extractions from a bulk aspen standard were included every twelve samples as process standards. These were used to correct for both between-batch and within-batch variation. For between-batch normalization, concentrations were scaled by dividing the overall mean of the first two process standard injections from each batch by the mean of the first two process standard injections within the given batch. Within each batch, process standards showed a linear decline over run time. To account for this drift, we fit a regression line to the mean concentration of the paired standards as a function of run order and applied the inverse of this function to all samples in the same batch. Concentrations of salicinoid phenolic glycosides (SPGs) were then quantified using standard curves, and values were converted to percent dry weight as: % d.w. = extraction volume*dilution factor*1.09*100*(sample conconcentration/(sample start weight (g)*1000)) where 1.09 is the empirically determined extraction efficiency.

#### CT Quantification

CTs were extracted in 70% acetone with 0.01% ascorbic acid and quantified using the acid-butanol colorimetric method(81). Extracts were mixed with 95:5 butanol:HCl containing ferric ammonium sulfate and incubated at 95°C for 40 minutes. Absorbance was measured at 550 nm. Procyanidin B2 (≥ 90% purity by HPLC, CAS 29106-49-8, MW 578.52 g/mol; Sigma-Aldrich, St. Louis, MO, USA, Cat. No. 42157) was used as the analytical standard.

#### Microbial Community Characterization Sampling and Processing

Microbial community sampling was conducted concurrently with chemical sampling. To minimize diel variation in microbiome composition, leaves were collected in the morning over 3–4 consecutive days. Equal numbers of samples were randomly selected from each plot daily.

Leaves were clipped from mid-canopy height into sterile tubes, washed in the field with 4-5 mL 0.1% Tween-20 solution, rinsed thoroughly with sterile deionized water, flash-frozen on dry ice, and stored at –80°C.

At field sites, sampling occurred from late July to early September 2022 and over five consecutive days in mid-July 2023. For each tree, 3–4 branches were collected using a 20-gauge shotgun. Four leaves per branch were clipped into sterile tubes, then washed and stored as above.

#### DNA Extraction and Sequencing Extraction

DNA and RNA were co-extracted from whole frozen leaves using a modified CTAB-based protocol (82–84). Leaves were ground in liquid nitrogen with sterile foil-lined mortars and pestles, then transferred to 2 mL microcentrifuge tubes. Cell lysis was performed in CTAB buffer (100 mM Tris-HCl pH 8.0, 1.4 M NaCl, 20 mM EDTA, 2% CTAB, 2% polyvinylpyrrolidone K30, 2% β-mercaptoethanol, 100 µg/mL Proteinase K) at 60°C for 15 minutes. Nucleic acids were extracted with 24:1 chloroform:isoamyl alcohol and purified through two rounds of extraction.

The final aqueous phase was split: 250 µL was set aside for RNA extraction. DNA was precipitated with NaCl, potassium acetate, and isopropanol, incubated at –20°C overnight, and centrifuged at 3300 rpm for 45 minutes. Pellets were washed with 80% ethanol, dried, and resuspended in nuclease-free water.

#### Amplicon Sequencing

The ITS1 region was amplified using a two-step PCR protocol using a set of frameshifting oligos (six frameshifted variants each, forward and reverse) modified from Regalado et al, (2020) (85). The first PCR was as described in Regalado et. al (6 μL 10X ThermoPol Taq buffer (NEB), 0.96 uL 5μM forward primer, 0.96 μL 5uM reverse primer, 1.2 μL 10mM dNTPs, 0.48 μL Taq DNA polymerase (NEB), nuclease free water to 55 μL and 5 μL DNA template. Blocking oligos were excluded. The reaction was then split across three 96-well PCR plates with 20 μL per reaction and run in triplicate under the following conditions: 94 C for 2 min; ten cycles of 94C for 30s, 55C for 30s, 72C for 30s; 72C for 3min. Amplified DNA was then pooled and cleaned with 1:1 SPRI beads prior to PCR2. The second PCR was set up as single 25 μL reactions per sample (5 μL Q5 PCR buffer (NEB), 0.0625 μL 100 μM universal forward primer, 1.25 μL 5 μM barcoded reverse primer, 0.5 μL 10 mM dNTPs, 0.25 μL Q5 polymerase (NEB), 4.875 μL nuclease-free water, and 13 μL template from PCR1). Thermal cycling conditions were: 94 °C for 1 min; 25 cycles of 94 °C for 20 s, 60 °C for 30 s, 72 °C for 30 s; and a final extension at 72 °C for 2 min. Amplicon size (∼400-450 bp) and adapter incorporation were confirmed by agarose gel electrophoresis. Amplicons were pooled by plate (10 μL/sample), cleaned with a 1:1.2 SPRI bead ratio and eluted in 300 μL molecular grade water. They were then concentrated (10x) at 40C on a vacuum centrifuge prior to size-selection (300–750 bp) on a BluePippin (1.5% gel). Libraries were sequenced on an Illumina MiSeq using a v3 600-cycle kit.

Reads were demultiplexed and adapters trimmed using scripts adapted from https://github.com/tkarasov/pathodopsis (86). Reads were filtered and denoised without merging using DADA2 (maxN=0; maxEE=2; truncQ=2; minLen=50) (87). Taxonomic assignment was performed on forward reads using the UNITE database v9.0 general FASTA release (88).

Assigned reads were further analyzed with phyloseq (v1.50.0). Taxonomic assignments were filtered to retain only fungal ASVs classified at the family level or lower. Samples with fewer than 500 total assigned reads were excluded from downstream analyses.

To filter for potential pathogens, we compiled a list from a literature search of families known or suspected to contain plant pathogens. We then filtered the taxa table to include only these families. In complement, we calculated the variance in relative abundances for each ASV and extracted a list of families with at least one ASV in the top quartile of variance in relative abundance. We then further filtered the taxa table by the list of high variance families.

## Statistical Analyses

Bray-Curtis dissimilarities were calculated from the full table of UPLC results, normalized by sample weight using the vegdist function in vegan (v2.6.10) ((89). Principal coordinates analysis (PCoA) was then performed on the resulting distance matrix with the pcoa function in ape (v5.8.1) ((90). Treatment groups were compared using permutational multivariate analysis of variance (PERMANOVA) with the adonis2 function in vegan.

To test for effects of drought treatment on specialized metabolites, we used linear mixed-effects models (lme4 package (v1.1.36) in R) ((91):

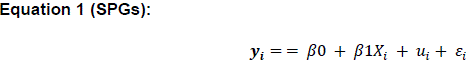

Where y_i_ is the metabolite concentration, X_i_ is drought treatment, u_i_ is the random intercept for genotype, and ɛ_i_ is the residual error.

Type III ANOVA (car package (v3.1.3)) was used to test for significant treatment effects (92). Post hoc pairwise comparisons were conducted using the emmeans package (v1.10.7) with Tukey correction for multiple testing (93).

Variance explained was calculated using the partR2 package(v0.9.2) ((94). Both Marginal and Conditional R² were calculated for each model. Marginal R² was used to estimate fixed effects (treatment), and the difference between conditional R² and marginal R² isolated the variance explained by genotype.

Log-fold changes in microbial taxa and clustering of co-varying ASVs were assessed using DESeq2 (v1.46.0) ((55). Comparisons of relative abundances between treatment groups were determined with the DESeq function using the Likelihood Ratio Test (“LRT”) with plot as a factor included as a covariate in the model(55).

To test whether group and chemistry explained variation in the relative abundance of individual taxa, we used linear mixed-effects models with plot specified as a random effect, implemented via the taxatree_models function in *microViz* (v0.12.7) ((95).

## References

1. T. J. Brodribb, J. Powers, H. Cochard, B. Choat, Hanging by a thread? Forests and drought. Science 368, 261–266 (2020).

2. W. M. Hammond, et al., Global field observations of tree die-off reveal hotter-drought fingerprint for Earth’s forests. Nat. Commun. 13, 1761 (2022).

3. A. Gazol, M. Pizarro, W. M. Hammond, C. D. Allen, J. J. Camarero, Droughts preceding tree mortality events have increased in duration and intensity, especially in dry biomes. Nat. Commun. 16, 5779 (2025).

4. A. T. Trugman, L. D. L. Anderegg, W. R. L. Anderegg, A. J. Das, N. L. Stephenson, Why is Tree Drought Mortality so Hard to Predict? Trends Ecol. Evol. 36, 520–532 (2021).

5. H. Hartmann, et al., Climate change risks to global forest health: Emergence of unexpected events of elevated tree mortality worldwide. Annu. Rev. Plant Biol. 73, 673–702 (2022).

6. W. R. L. Anderegg, et al., Tree mortality from drought, insects, and their interactions in a changing climate. New Phytol. 208, 674–683 (2015).

7. H. Hartmann, et al., Research frontiers for improving our understanding of drought-induced tree and forest mortality. New Phytol. 218, 15–28 (2018).

8. C. Gely, S. G. W. Laurance, N. E. Stork, How do herbivorous insects respond to drought stress in trees? Biol. Rev. Camb. Philos. Soc. 95, 434–448 (2020).

9. T. E. Kolb, et al., Observed and anticipated impacts of drought on forest insects and diseases in the United States. For. Ecol. Manage. 380, 321–334 (2016).

10. H. Jactel, et al., Drought effects on damage by forest insects and pathogens: a meta-analysis. Glob. Chang. Biol. 18, 267–276 (2012).

11. M. L. Gaylord, et al., Drought predisposes piñon–juniper woodlands to insect attacks and mortality. New Phytol. 198, 567–578 (2013).

12. J. Huang, et al., Tree defence and bark beetles in a drying world: carbon partitioning, functioning and modelling. New Phytol. 225, 26–36 (2020).

13. R. Nakabayashi, K. Saito, Integrated metabolomics for abiotic stress responses in plants. Curr. Opin. Plant Biol. 24, 10–16 (2015).

14. P. Trivedi, J. E. Leach, S. G. Tringe, T. Sa, B. K. Singh, Plant-microbiome interactions: from community assembly to plant health. Nat. Rev. Microbiol. 18, 607–621 (2020).

15. A. E. Burns, R. M. Gleadow, I. E. Woodrow, Light alters the allocation of nitrogen to cyanogenic glycosides in Eucalyptus cladocalyx. Oecologia 133, 288–294 (2002).

16. R. M. Gleadow, W. J. Foley, I. E. Woodrow, Enhanced CO2 alters the relationship between photosynthesis and defence in cyanogenic Eucalyptus cladocalyx F. Muell. Plant Cell Environ. 21, 12–22 (1998).

17. J. Huang, et al., Eyes on the future – evidence for trade-offs between growth, storage and defense in Norway spruce. New Phytol. 222, 144–158 (2019).

18. O. L. Cope, R. L. Lindroth, A. Helm, K. Keefover-Ring, E. L. Kruger, Trait plasticity and trade-offs shape intra-specific variation in competitive response in a foundation tree species. New Phytol. 230, 710–719 (2021).

19. C. T. Cole, M. T. Stevens, J. E. Anderson, R. L. Lindroth, Heterozygosity, gender, and the growth-defense trade-off in quaking aspen. Oecologia 181, 381–390 (2016).

20. S. Munné-Bosch, L. Alegre, Changes in carotenoids, tocopherols and diterpenes during drought and recovery, and the biological significance of chlorophyll loss in Rosmarinus officinalis plants. Planta 210, 925–931 (2000).

21. M. Moroldo, et al., Genetic control of abiotic stress-related specialized metabolites in sunflower. BMC Genomics 25, 199 (2024).

22. M. C. Cambon, et al., Drought tolerance traits in Neotropical trees correlate with the composition of phyllosphere fungal communities. Phytobiomes J. (2022). 10.1094/pbiomes-04-22-0023-r.

23. B. K. Singh, et al., Climate change impacts on plant pathogens, food security and paths forward. Nat. Rev. Microbiol. (2023). 10.1038/s41579-023-00900-7.

24. D. L. Forrister, M.-J. Endara, G. C. Younkin, P. D. Coley, T. A. Kursar, Herbivores as drivers of negative density dependence in tropical forest saplings. Science 363, 1213–1216 (2019).

25. A. Piasecka, et al., Drought-related secondary metabolites of barley (Hordeum vulgare L.) leaves and their metabolomic quantitative trait loci. Plant J. 89, 898–913 (2017).

26. M. Shamloo, et al., Effects of genotype and temperature on accumulation of plant secondary metabolites in Canadian and Australian wheat grown under controlled environments. Sci. Rep. 7, 9133 (2017).

27. U. Sarker, S. Oba, Drought stress effects on growth, ROS markers, compatible solutes, phenolics, flavonoids, and antioxidant activity in Amaranthus tricolor. Appl. Biochem. Biotechnol. 186, 999–1016 (2018).

28. A. Khaleghi, et al., Morphological, physiochemical and antioxidant responses of Maclura pomifera to drought stress. Sci. Rep. 9, 19250 (2019).

29. J. Koricheva, S. Larsson, E. Haukioja, M. Keinänen, M. Keinanen, Regulation of woody plant secondary metabolism by resource availability: Hypothesis testing by means of meta-analysis. Oikos 83, 212 (1998).

30. J. K. Holopainen, et al., Climate change effects on secondary compounds of forest trees in the northern hemisphere. Front. Plant Sci. 9, 1445 (2018).

31. I. Nogués, et al., Physiological and antioxidant responses of*Quercus ilex*to drought in two different seasons. Plant Biosyst. 148, 268–278 (2014).

32. A. Rivas-Ubach, et al., Drought enhances folivory by shifting foliar metabolomes in Quercus ilex trees. New Phytologist 202, 874–885 (2014).

33. E. A. Gerson, R. G. Kelsey, Piperidine alkaloids in North American Pinus taxa: implications for chemosystematics. Biochem. Syst. Ecol. 32, 63–74 (2004).

34. A. Kleiber, et al., Drought effects on root and needle terpenoid content of a coastal and an interior Douglas fir provenance. Tree Physiol. 37, 1648–1658 (2017).

35. S. C. Malone, et al., Water, not carbon, drives drought-constraints on stem terpene defense against simulated bark beetle attack in Pinus edulis. New Phytol. 245, 318–331 (2025).

36. R. A. Thompson, et al., Local carbon reserves are insufficient for phloem terpene induction during drought in Pinus edulis in response to bark beetle-associated fungi. New Phytol. 244, 654–669 (2024).

37. J. Laoué, et al., A decade of rain exclusion in a Mediterranean forest reveals trade-offs of leaf chemical defenses and drought legacy effects. Sci. Rep. 14, 24119 (2024).

38. M. Eisenring, et al., Legacy effects of premature defoliation in response to an extreme drought event modulate phytochemical profiles with subtle consequences for leaf herbivory in European beech. New Phytol. 242, 2495–2509 (2024).

39. Y.-G. Zhu, et al., Impacts of global change on the phyllosphere microbiome. New Phytol. 234, 1977–1986 (2022).

40. T. L. Karasov, et al., Continental-scale associations of Arabidopsis thaliana phyllosphere members with host genotype and drought. Nat. Microbiol. 9, 2748–2758 (2024).

41. Q.-L. Chen, et al., Precipitation increases the abundance of fungal plant pathogens in Eucalyptus phyllosphere. Environ. Microbiol. 23, 7688–7700 (2021).

42. J. Peñuelas, L. Rico, R. Ogaya, A. S. Jump, J. Terradas, Summer season and long-term drought increase the richness of bacteria and fungi in the foliar phyllosphere of Quercus ilex in a mixed Mediterranean forest. Plant Biol. 14, 565–575 (2012).

43. L. Rico, R. Ogaya, J. Terradas, J. Peñuelas, Community structures of N2 –fixing bacteria associated with the phyllosphere of a Holm oak forest and their response to drought. Plant Biol. 16, 586–593 (2014).

44. T. Lin, et al., Drought stress-mediated differences in phyllosphere microbiome and associated pathogen resistance between male and female poplars. Plant J. 115, 1100–1113 (2023).

45. R. Khlifa, et al., Conifer epiphytic phyllosphere bacterial communities respond more strongly to rain exclusion and host species identity than to soil water content. For. Ecol. Manage. 581, 122554 (2025).

46. A. C. H. Jaeger, M. Hartmann, R. F. Conz, J. Six, E. F. Solly, Prolonged water limitation shifts the soil microbiome from copiotrophic to oligotrophic lifestyles in Scots pine mesocosms. Environ. Microbiol. Rep. 16, e13211 (2024).

47. A. J. M. Hopkins, et al., Drought legacy interacts with wildfire to alter soil microbial communities in a Mediterranean climate-type forest. Sci. Total Environ. 915, 170111 (2024).

48. K. F. Rubert-Nason, R. L. Lindroth, Causes and consequences of condensed tannin variation in Populus. Recent Advances in Polyphenol Research [Preprint] (2021). Available at: https://onlinelibrary.wiley.com/doi/10.1002/9781119545958.ch4.

49. J. D. C. Hemming, R. L. Lindroth, Intraspecific variation in aspen phytochemistry: effects on performance of gypsy moths and forest tent caterpillars. Oecologia 103, 79–88 (1995).

50. A. Gruppe, M. Fußeder, R. Schopf, Short rotation plantations of aspen and balsam poplar on former arable land in Germany: defoliating insects and leaf constituents. For. Ecol. Manage. 121, 113–122 (1999).

51. C. Ullah, et al., Flavan-3-ols are an effective chemical defense against rust infection. Plant Physiol. 175, 1560–1578 (2017).

52. M. V. Van Nuland, et al., Above– and belowground fungal biodiversity of Populus trees on a continental scale. Nat. Microbiol. 8, 2406–2419 (2023).

53. K. L. Kerr, J. C. Fickle, W. R. L. Anderegg, Decoupling of functional traits from intraspecific patterns of growth and drought stress resistance. New Phytol. 239, 174–188 (2023).

54. J. C. Fickle, G. Vargas G, W. R. L. Anderegg, Ring-specific vulnerability to embolism reveals accumulation of damage in the xylem. New Phytol. 246, 2046–2058 (2025).

55. M. I. Love, W. Huber, S. Anders, Moderated estimation of fold change and dispersion for RNA-seq data with DESeq2. Genome Biol. 15, 550 (2014).

56. J. R. Donaldson, M. T. Stevens, H. R. Barnhill, R. L. Lindroth, Age-related shifts in leaf chemistry of clonal aspen (Populus tremuloides). J. Chem. Ecol. 32, 1415–1429 (2006).

57. R. L. Lindroth, T. L. Osier, H. R. H. Barnhill, S. A. Wood, Effects of genotype and nutrient availability on phytochemistry of trembling aspen (Populus tremuloides Michx.) during leaf senescence. Biochem. Syst. Ecol. 30, 297–307 (2002).

58. J. D. C. Hemming, R. L. Lindroth, Effects of Light and Nutrient Availability on Aspen. J. Chem. Ecol. 25, 1687–1714 (1999).

59. J. R. Donaldson, R. L. Lindroth, Genetics, environment, and their interaction determine efficacy of chemical defense in trembling aspen. Ecology 88, 729–739 (2007).

60. R. L. Lindroth, et al., Phenotypic variation in phytochemical defense of trembling Aspen in western North America: Genetics, development, and geography. J. Chem. Ecol. 49, 235–250 (2023).

61. G. A. Boeckler, et al., Transgenic upregulation of the condensed tannin pathway in poplar leads to a dramatic shift in leaf palatability for two tree-feeding Lepidoptera. J. Chem. Ecol. 40, 150–158 (2014).

62. G. Gourlay, B. J. Hawkins, A. Albert, J.-P. Schnitzler, C. Peter Constabel, Condensed tannins as antioxidants that protect poplar against oxidative stress from drought and UV-B. Plant Cell Environ. 45, 362–377 (2022).

63. G. Gourlay, C. P. Constabel, Condensed tannins are inducible antioxidants and protect hybrid poplar against oxidative stress. Tree Physiol. 39, 345–355 (2019).

64. J. J. Kelly, et al., Alteration of microbial communities colonizing leaf litter in a temperate woodland stream by growth of trees under conditions of elevated atmospheric CO2. Appl. Environ. Microbiol. 76, 4950–4959 (2010).

65. R. S. Winder, J. Lamarche, C. P. Constabel, R. C. Hamelin, The effects of high-tannin leaf litter from transgenic poplars on microbial communities in microcosm soils. Front. Microbiol. 4, 290 (2013).

66. K. F. Rubert-Nason, J. J. Couture, E. A. Gryzmala, P. A. Townsend, R. L. Lindroth, Vernal freeze damage and genetic variation alter tree growth, chemistry, and insect interactions. Plant Cell Environ. 40, 2743–2753 (2017).

67. R. L. Lindroth, J. D. C. Hemming, Responses of the Gypsy Moth (Lepidoptera: Lymantriidae) to Tremulacin, an Aspen Phenolic Glycoside. Environ. Entomol. 19, 842–847 (1990).

68. M. A. Falk, R. L. Lindroth, K. Keefover-Ring, K. F. Raffa, Genetic variation in aspen phytochemical patterns structures windows of opportunity for gypsy moth larvae. Oecologia 187, 471–482 (2018).

69. M. M. Dudley, N. A. Tisserat, W. R. Jacobi, J. Negrón, J. E. Stewart, Pathogenicity and distribution of two species of Cytospora on Populus tremuloides in portions of the Rocky Mountains and midwest in the United States. For. Ecol. Manage. 468, 118168 (2020).

70. S. B. Marchetti, J. J. Worrall, T. Eager, Secondary insects and diseases contribute to sudden aspen decline in southwestern Colorado, USA. Can. J. For. Res. 41, 2315–2325 (2011).

71. B. R. Albrectsen, et al., Endophytic fungi in European aspen (Populus tremula) leaves—diversity, detection, and a suggested correlation with herbivory resistance. Fungal Divers. 41, 17–28 (2010).

72. A. B. Siddique, S. Parveen, M. Z. Rahman, J. Rahman, Revisiting plant stress memory: mechanisms and contribution to stress adaptation. Physiol. Mol. Biol. Plants 30, 349–367 (2024).

73. P. A. Auler, et al., Stress memory of physiological, biochemical and metabolomic responses in two different rice genotypes under drought stress: The scale matters. Plant Science [Preprint] (2021). Available at: 10.1016/j.plantsci.2021.110994.

74. M. Sharma, et al., Understanding plant stress memory response for abiotic stress resilience: Molecular insights and prospects. Plant Physiol. Biochem. 179, 10–24 (2022).

75. J. J. Worrall, et al., Effects and etiology of sudden aspen decline in southwestern Colorado, USA. For. Ecol. Manage. 260, 638–648 (2010).

76. J. J. Worrall, A. G. Keck, S. B. Marchetti, Populus tremuloides stands continue to deteriorate after drought-incited sudden aspen decline. Can. J. For. Res. (2015).

77. A. Dai, Increasing drought under global warming in observations and models. Nat. Clim. Chang. 3, 52–58 (2013).

78. B. I. Cook, T. R. Ault, J. E. Smerdon, Unprecedented 21st century drought risk in the American Southwest and Central Plains. Sci. Adv. 1, e1400082 (2015).

79. K. Rubert-Nason, K. Keefover-Ring, R. L. Lindroth, Purification and Analysis of Salicinoids. Curr. Anal. Chem. 14, 423–429 (2018).

80. R. Schmid, et al., Integrative analysis of multimodal mass spectrometry data in MZmine 3. Nat. Biotechnol. 41, 447–449 (2023).

81. L. J. Porter, L. N. Hrstich, B. G. Chan, The conversion of procyanidins and prodelphinidins to cyanidin and delphinidin. Phytochemistry 25, 223–230 (1985).

82. S. Chang, J. Puryear, J. Cairney, A simple and efficient method for isolating RNA from pine trees. Plant Mol. Biol. Rep. 11, 113–116 (1993).

83. S. Sasi, et al., DNA-free high-quality RNA extraction from 39 difficult-to-extract plant species (representing seasonal tissues and tissue types) of 32 families, and its validation for downstream molecular applications. Plant Methods 19, 84 (2023).

84. D. S. Lundberg, et al., A major trade-off between growth and defense in Arabidopsis thaliana can vanish in field conditions. PLoS Biol. 23, e3003237 (2025).

85. J. Regalado, D. S. Lundberg, O. Deusch, S. Kersten, Combining whole-genome shotgun sequencing and rRNA gene amplicon analyses to improve detection of microbe–microbe interaction networks in plant …. ISME J. (2020).

86. T. Karasov, tkarasov/Pathodopsis. [Preprint] (2023). Available at: https://github.com/tkarasov/pathodopsis.

87. B. J. Callahan, et al., DADA2: High-resolution sample inference from Illumina amplicon data. Nat. Methods 13, 581–583 (2016).

88. K. Abarenkov, et al., The UNITE database for molecular identification and taxonomic communication of fungi and other eukaryotes: sequences, taxa and classifications reconsidered. Nucleic Acids Res. 52, D791–D797 (2024).

89. J. Oksanen, et al., vegan: Community Ecology Package. [Preprint] (2025). Available at: https://vegandevs.github.io/vegan/.

90. E. Paradis, J. Claude, K. Strimmer, APE: Analyses of Phylogenetics and Evolution in R language. Bioinformatics 20, 289–290 (2004).

91. D. Bates, M. Mächler, B. Bolker, S. Walker, Fitting linear mixed-effects models Usinglme4. J. Stat. Softw. 67, 1–48 (2015).

92. J. Fox, S. Weisberg, An R Companion to Applied Regression. [Preprint] (2019). Available at: https://www.john-fox.ca/Companion/.

93. R. V. Lenth, emmeans: Estimated Marginal Means, aka Least-Squares Means. [Preprint] (2025). Available at: https://rvlenth.github.io/emmeans/.

94. M. A. Stoffel, S. Nakagawa, H. Schielzeth, partR2: Partitioning R2 in generalized linear mixed models. PeerJ [Preprint] (2021). Available at: 10.7717/peerj.11414.

95. D. J. M. Barnett, I. C. W. Arts, J. Penders, microViz: an R package for microbiome data visualization and statistics. Journal of Open Source Software [Preprint] (2021). Available at: 10.21105/joss.03201.

