## supplementary_figures for "Functional memory of drought affects leaf chemical defenses and microbial interactions in aspen"

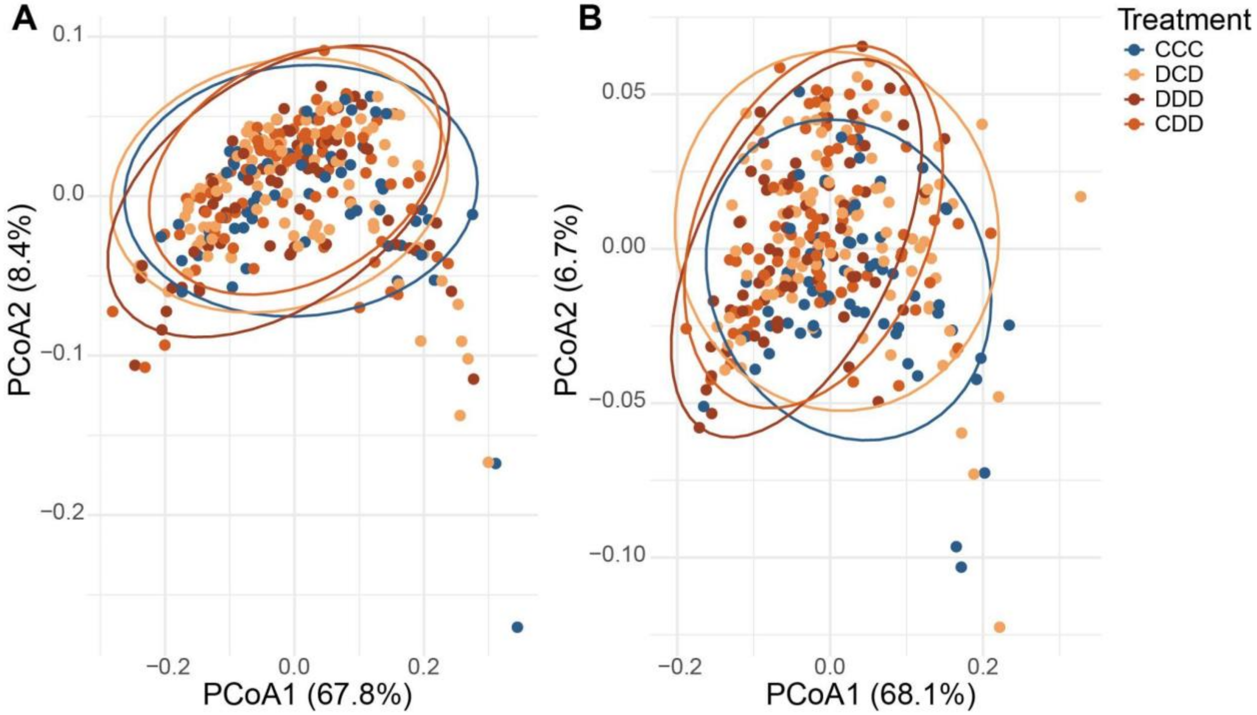


Supplementary Figure 1. **Total foliar chemistry separates by drought treatment.** PCoA ordination for all UPLC peaks normalized by sample starting weight. Similar variance in samples was explained by each of the first two PCoA axes in 2022 (A) and 2023(B).

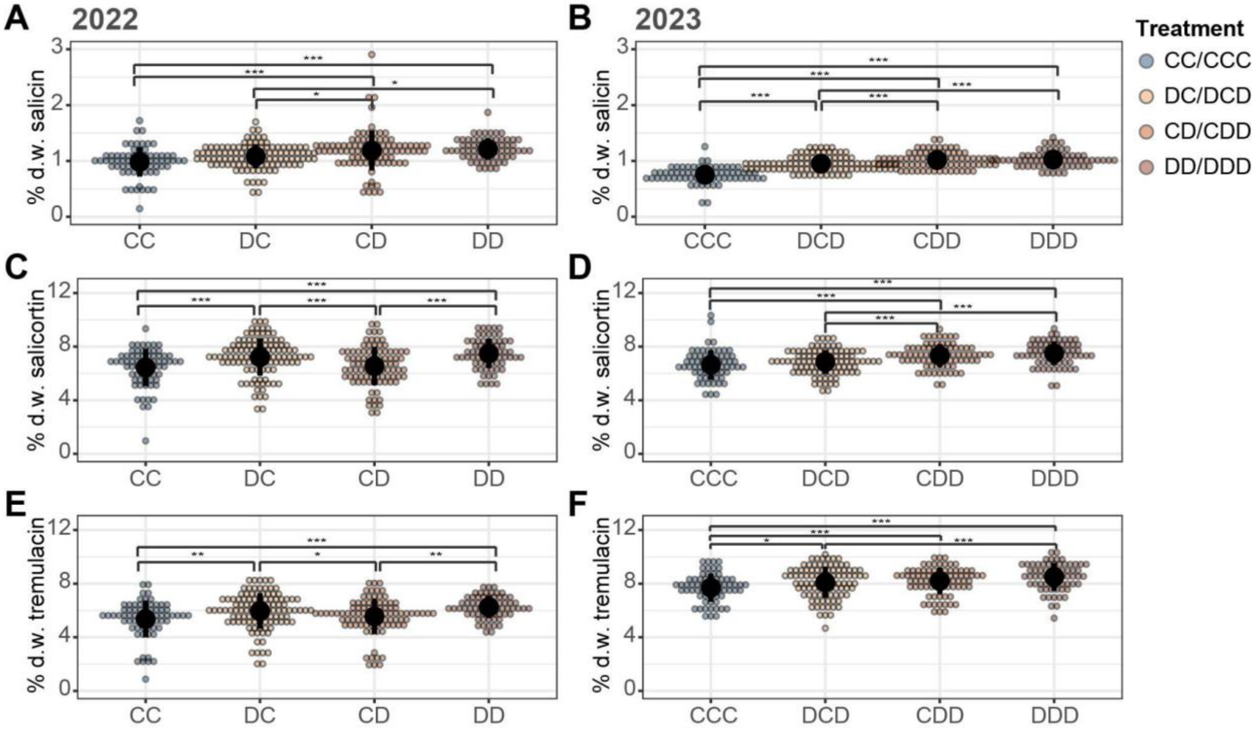

Supplementary Figure 2. **Drought memory effects on allocation to individual SPGs** Percent dry weight (% d.w.) allocation to salicin (A-B), salicortin (C-D) and tremulacin(E-F) in 2022 (A,C,E) and 2023 (B,D,F). Colored dots represent individual samples; black lines show means ± SE. Pairwise comparisons were performed using *emmeans* with Tukey-adjusted ps: (***) < 0.001, (**) <0.01, (*) <0.05. Sample sizes: n ≤ 51 (CCC, DDD) or n ≤ 74 (CDD, DCD).

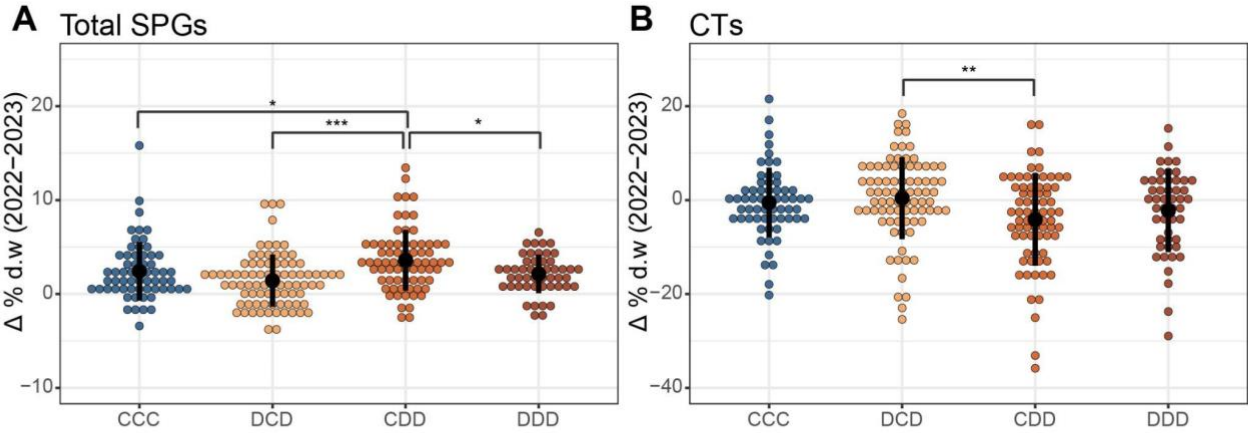


Supplementary Figure 3 **Changing allocations to defensive chemistry differ by treatment group.** Difference in percent dry weight (Δ % d.w.) from 2022 to 2023 in SPGs (A) and CTs (B). Colored dots represent individual samples; black lines show means ± SE. Pairwise comparisons were performed using *emmeans* with Tukey-adjusted p-value: (***) < 0.001, (**) <0.01, (*) <0.05. Sample sizes: n ≤ 51 (CCC, DDD) or n ≤ 74 (CDD, DCD).

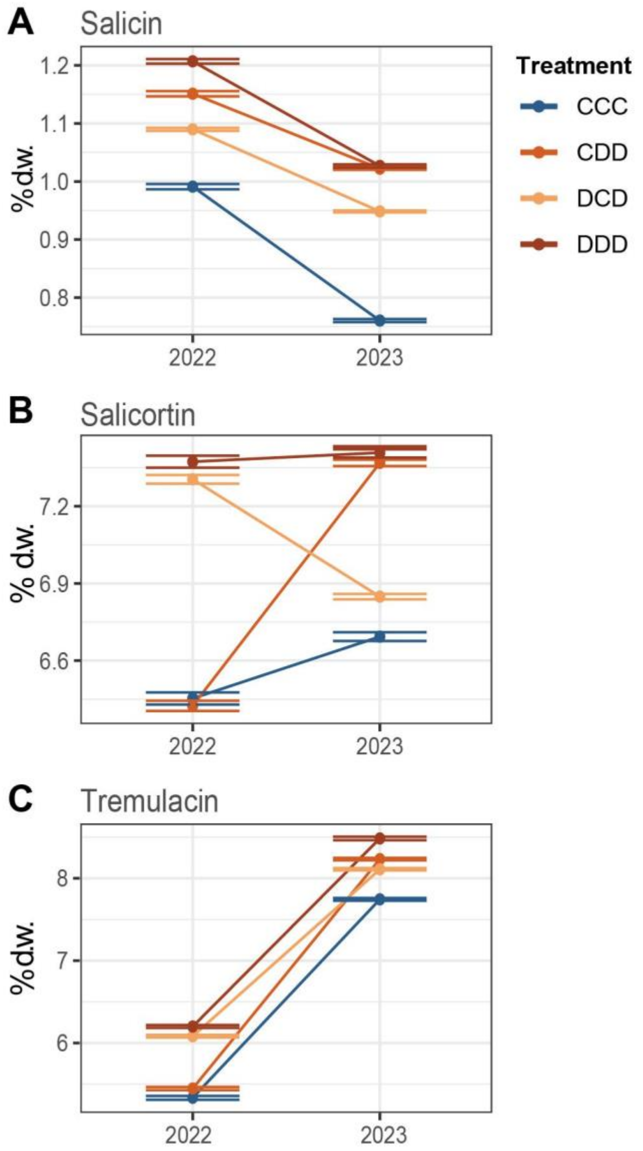


Supplementary figure 4. **Change in individual SPGs from 2022 to 2023**. Mean total percent dry weight (% d.w.) ± SE for salicin (A), salicortin (B) and tremulacin (C) in 2022 and 2023 by treatment group.

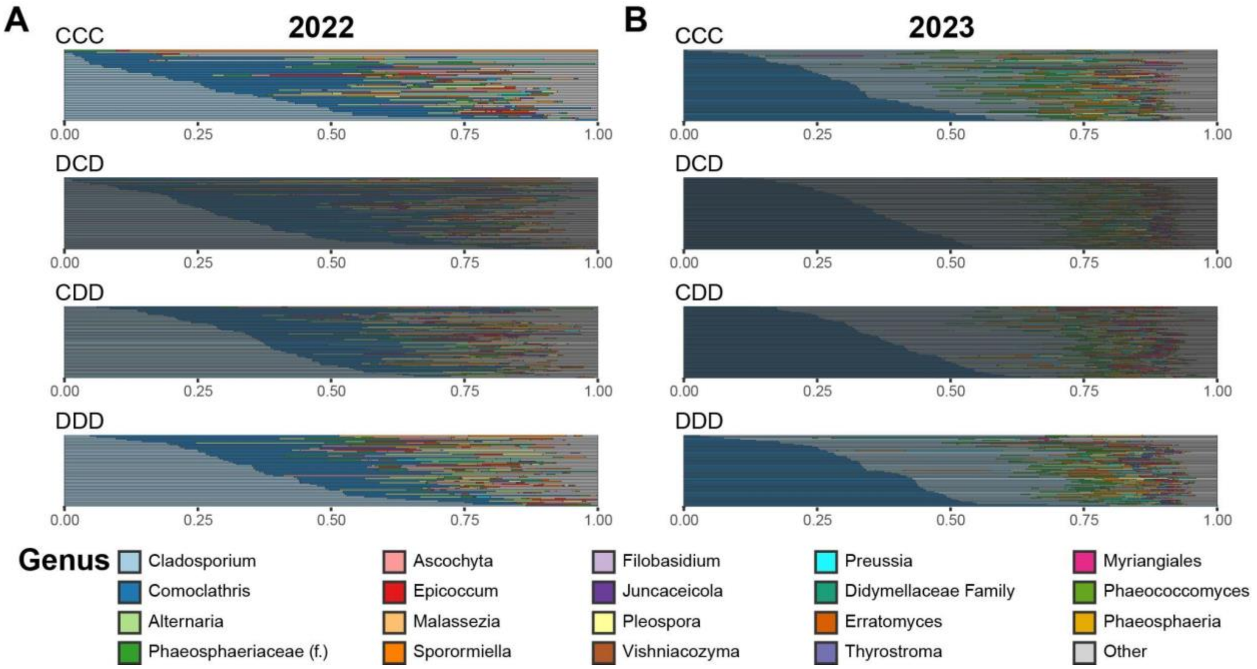


Supplementary figure 5. **Relative abundances of fungal genera across treatment groups.** Samples collected from the common garden at peak drought in 2022 and 2023. ITS1 amplified from single leaves collected in the common garden. Relative abundances of fungal genera trees in 2022 (A) and 2023 (B) by treatment group.
